# The *Drosophila* Tumour Suppressor Lgl and Vap33 activate the Hippo pathway by a dual mechanism, involving RtGEF/Git/Arf79F and inhibition of the V-ATPase

**DOI:** 10.1101/2023.07.10.548302

**Authors:** Marta Portela, Swastik Mukherjee, Sayantanee Paul, John E. La Marca, Linda M. Parsons, Alexey Veraksa, Helena E. Richardson

## Abstract

The tumour suppressor, Lethal (2) giant larvae (Lgl), is an evolutionarily conserved protein that was discovered in the vinegar fly, *Drosophila*, where its depletion results in tissue overgrowth and loss of cell polarity and tissue architecture. Our previous studies have revealed a new role for Lgl in linking cell polarity and tissue growth through regulation of the Notch (proliferation and differentiation) and the Hippo (negative tissue growth control) signalling pathways. Moreover, Lgl regulates vesicle acidification, via the Vacuolar ATPase (V-ATPase), and we showed that Lgl inhibits V-ATPase activity through Vap33 (a Vamp (v-SNARE)-associated protein, involved in endo-lysosomal trafficking) to regulate the Notch pathway. However, how Lgl acts to regulate the Hippo pathway was unclear. In this current study, we show that V-ATPase activity inhibits the Hippo pathway, whereas Vap33 acts to activate Hippo signalling. Using an *in vivo* affinity-purification approach we found that Vap33 binds to the actin cytoskeletal regulators RtGEF (Pix, a Rho-type guanine nucleotide exchange factor) and Git (G protein-coupled receptor kinase interacting ArfGAP), which also bind to the Hpo protein kinase, and are involved in the activation of the Hippo pathway. Vap33 genetically interacts with RtGEF and Git in Hippo pathway regulation. Additionally, we show that the ADP ribosylation factor Arf79F (Arf1), which is a Hpo interactor, is involved in the inhibition of the Hippo pathway. Altogether our data suggests that Lgl acts via Vap33 to activate the Hippo pathway by a dual mechanism, 1) through interaction with RtGEF/Git/Arf79F, and 2) through interaction and inhibition of the V-ATPase, thereby controlling epithelial tissue growth.

## Introduction

Deregulation of cell polarity and the epithelial-to mesenchymal transition are hallmarks of human cancer [1-3]. Key regulators of cell polarity, Lgl, Scribble (Scrib), and Discs large (Dlg), were discovered in *Drosophila* as neoplastic tumour suppressors; these proteins regulate apical-basal cell polarity via their antagonistic interactions with the Par-aPKC and Crb polarity complexes, and also limit cell proliferation [3]. Lgl functions to antagonise the activity of aPKC (atypical protein kinase C), and conversely aPKC phosphorylates and inhibits Lgl, in cell polarity regulation and tissue growth control [3]. In *Drosophila*, Lgl-aPKC play a role in the control of tissue growth (cell proliferation and survival) that is distinct from their role in cell polarity regulation; *lgl* mutation or aPKC-activation in eye epithelial tissue exhibits increased cell proliferation and survival without loss of apico-basal cell polarity [4, 5]. Thus, the Lgl-aPKC axis regulates tissue growth independent of cell polarity effects.

The human Lgl ortholog, LLGL1, has a conserved function with *Drosophila* Lgl, as its expression in *Drosophila* rescues the tumourigenic defects of *lgl* mutants [5, 6]. Reduced expression or mutations in LLGL1 (HUGL1), are associated with hepatocellular carcinoma [7], malignant melanoma [8] and colorectal cancer [9]. Similarly, aberrant localization or deletion of the second human Lgl homolog, LLGL2 (HUGL2), is associated with gastric epithelial dysplasia and adenocarcinoma [10], and with pancreatic intraepithelial neoplasia and ductal adenocarcinoma [11]. Mislocalization of LLGL1/2 is also observed in other human cancers, including lung adenocarcinoma [12] and ovarian cancer [13]. As occurs in *Drosophila*, the mislocalization/dysfunction of LLGL1/2 in cancer is also associated with altered aPKC localization/activity [12, 13].

We made the novel discovery that in *Drosophila* Lgl acts independent of its apico-basal cell polarity role to regulate the Salvador-Warts-Hippo (Hippo) negative tissue growth control pathway [4, 5]. The core of the Hippo pathway involves the serine-threonine protein kinases, Hippo (Hpo) and Warts (Wts), which respond to cell-cell contact and tissue architectural cues to control tissue growth via phosphorylating the co-transcriptional activator, Yorkie (Yki), which regulates cell proliferation genes (e.g. Cyclin E) and cell survival genes (e.g. Diap1) [14, 15]. Lgl depletion or aPKC activation (which inhibits Lgl) impairs the Hippo pathway via mislocalization of the Hpo protein, away from the apical cortex, where it is normally activated by apical cues [5, 16]. Thus, Lgl-aPKC controls the Hippo pathway by regulating Hpo localization and activity. In mammalian systems, deregulation of Lgl/aPKC impairs Hippo signalling and induces cell transformation, as aPKC’s association with the Hpo orthologs, MST1/2, and uncoupling MST from the downstream kinase, LATS (Wts), thereby leading to increased nuclear YAP (Yki) activity [17], consistent with what we observe in *Drosophila* [5].

We have also shown that in *Drosophila* Lgl plays a novel regulatory role in ligand-dependent Notch signalling [18, 19]. Notch activation depends on the cleavage by Adam proteases to produce Notch^ext^, and then processing by γ-secretase to produce the active form, Notch^ICD^, which translocates to the nucleus to activate transcription of target genes, such as the *E(spl)* complex, as well as the cell proliferation and survival genes [20]. We found that intracellular Notch accumulated in *lgl* mutant tissue, resulting in elevated Notch signalling that contributes to the overgrowth defects in *lgl* mutant tissue [5, 18, 19]. In mouse neural development, *Lgl1* knockout exhibits elevated Notch signalling, which is associated with hyperproliferation and differentiation defects [21], whilst in zebrafish, *Lgl1* knockdown in the developing retina leads to elevated Notch signalling and neurogenesis defects, which are rescued by blocking Notch activity [22].

Intriguingly, we recently found that *lgl* mutant tissue exhibited increased LysoTracker incorporation [18, 19], indicating that there is elevated vesicle acidification, due to the activity of the Vacuolar-ATPase (V-ATPase). In the Notch signalling pathway, γ-secretase activity is dependent on vesicle acidification regulated by V-ATPase activity [23, 24]. Therefore, in *lgl* mutant tissue the increased V-ATPase activity elevates γ-secretase activity and Notch cleavage, forming Notch^ICD^, and leading to elevated Notch target gene expression. Consistent with this, genetically or chemically reducing vesicle acidification or V-ATPase function, reduced the elevated Notch signalling in *lgl* mutant tissue [18, 19]. Thus, elevated Notch signalling in *lgl* mutant tissue is due to increased vesicle acidification and elevated γ-secretase activity.

To identify novel proteins that link Lgl to the V-ATPase, we undertook affinity purification-mass spectrometry (AP-MS) analysis of Lgl in *Drosophila* S2 tissue culture cells [25]. Lgl interacted with aPKC and Par6 with high significance as expected. Among the novel interactors of Lgl that bound at high significance, the standout protein involved in endocytosis was the VAMP-(v-SNARE)-associated protein, Vap33, which is an ortholog of human VAPA/VAPB [25]. Vap33 (VAPA/B) physically and genetically interacts with endocytic regulators and is involved in endo-lysosomal trafficking, with mutations in *Drosophila Vap33* and human *VAPB* resulting in endocytic defects, including the accumulation of the early endosome Rab5 marker [26], a phenotype we also observed in *Drosophila lgl* mutant tissue [18, 19]. We confirmed the binding of Vap33 to Lgl by co-immunoprecipitations (co-IPs) from S2 cells and *in vivo* in *Drosophila* epithelial cells [25] by using proximity ligation assays (PLA) [27]. Comparison of our data with the global *Drosophila* proteomics network revealed that Vap33 and Lgl form a network with V-ATPase subunit proteins [25, 28]. Interaction with human VAPA/B and V-ATPase proteins was also evident in the human proteome [25, 29]. We also found that *Vap33* overexpression rescued the elevated Notch signalling in *lgl* mutant eye epithelial clones and the *lgl* mutant adult eye phenotype, whilst knockdown of *Vap33* enhanced these eye defects [25]. *Vap33* overexpression also reduced V-ATPase activity, as assayed by LysoTracker levels [25, 30]. Moreover, in *lgl*-knockdown S2 cells or in *lgl* mutant tissue the interaction between Vap33 and V-ATPase components was decreased [25]. Thus, Lgl binds to and facilitates the binding of Vap33 to V-ATPase components, which inhibits V-ATPase activity, thereby controlling vesicle acidity, γ-secretase activity and Notch signalling.

Whilst our previous studies have dissected how Lgl regulates the Notch pathway, it is currently not known precisely how Lgl regulates the Hippo pathway. Thus, in this study, we focused on the mechanism of this regulation. We show that the V-ATPase inhibits the Hippo pathway and conversely Vap33 activates the Hippo pathway. Mechanistically, Vap33 is connected to the Hippo pathway by interacting with the cytoskeletal regulators, RtGEF (Pix), Git, and Arf79F (Arf1), which bind to the Hpo protein kinase. Our findings are consistent with a model whereby Lgl-Vap33 promote Hippo signalling via a dual mechanism through interaction with RtGEF-Git-Arf79F and by inhibiting the V-ATPase.

## Results

### The Hippo signalling pathway is negatively regulated by V-ATPase activity in *Drosophila*

Our previous studies have revealed the involvement of Lgl in the negative regulation of the V-ATPase [25], which is an important regulator of the Notch pathway [23, 24]. Since the *lgl* mutant adult eye phenotype (which is due to both Notch and Hippo pathway deregulation) was suppressed by reducing V-ATPase levels (more so than individually inhibiting Notch signalling or reducing Yki/Sd activity [5, 18, 19]), we suspected that the V-ATPase activity might be involved in the regulation of the Hippo pathway. Indeed, reducing V-ATPase levels/activity using a *Vha68-2* RNAi line resulted in higher Hippo pathway activity as assessed by reduced Yki nuclear staining, and lower expression of the Yki target, *expanded (ex)-lacZ* (**Fig 1B**) compared to the *wild-type* control (**Fig 1A**, quantified in **Fig 1E**). Conversely, knockdown of the core Hippo pathway gene, *wts*, resulted in impaired Hippo pathway signalling, as assessed by increased Yki nuclear staining and Yki target gene expression (*ex-lacZ*) (**Fig 1C)**, and a similar level of pathway impairment was observed upon activating the V-ATPase by overexpressing *Vha44* [30] (**Fig 1D**, quantified in **Fig 1E**). Thus, the V-ATPase negatively regulates the Hippo signalling pathway, leading to the upregulation of Yki activity.

**Fig 1:**
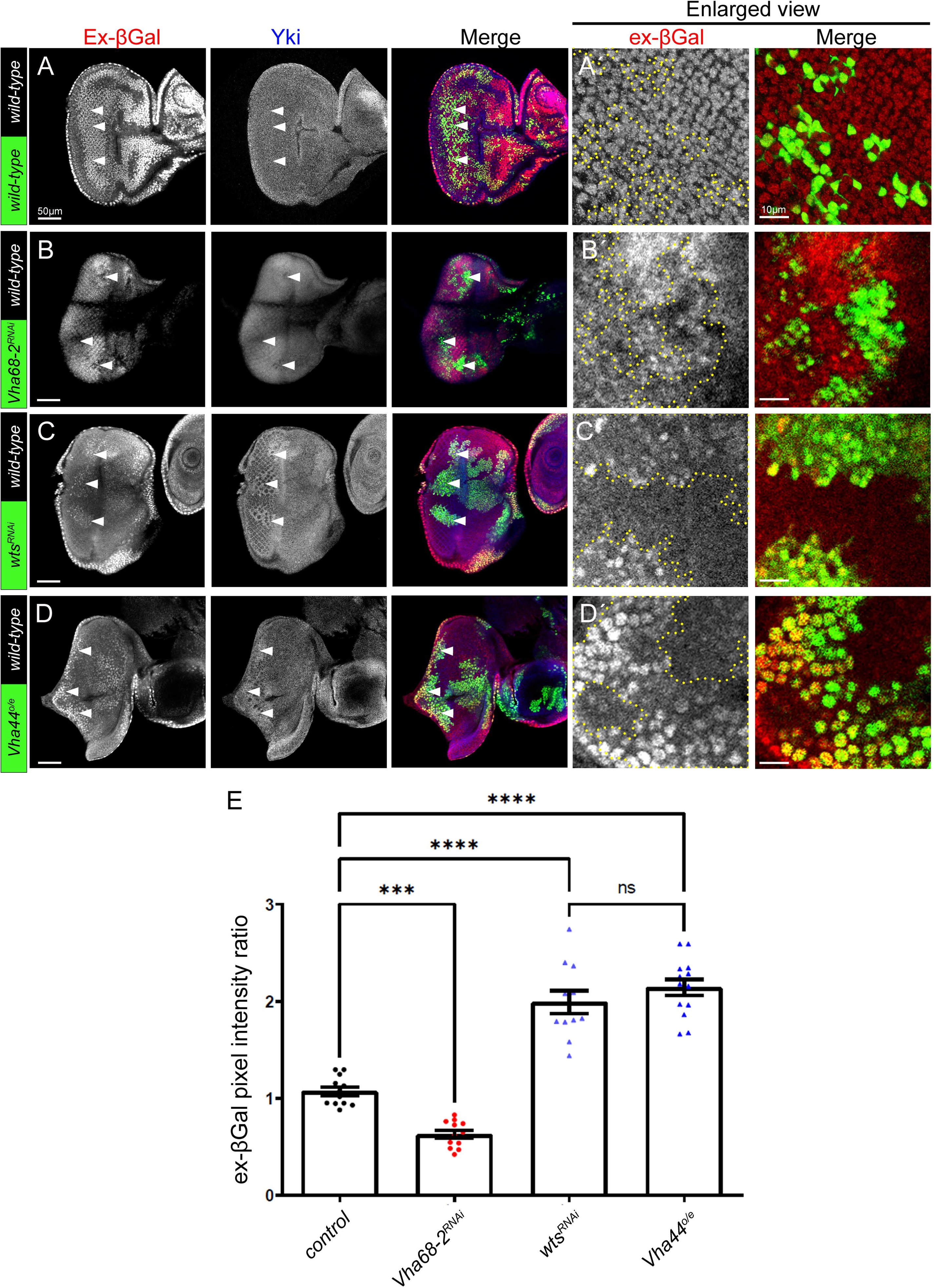
The Hippo signalling pathway is negatively regulated by V-ATPase activity in *Drosophila*. (A-D) Confocal planar sections of mosaic eye discs containing the Yki target reporter, *ex-lacZ*, stained for βGal (grey, or red in merges), and for Yki (grey, or blue in merges). Example GFP-positive clones are indicated by arrowheads. (A) Example of a control mosaic disc showing endogenous expression of *ex-lacZ* and Yki in the eye epithelium. (B) Example of a *Vha68-2^RNAi^* mosaic disc, with RNAi-expressing clones being GFP-positive. *Vha68-2* knockdown leads to downregulation of the Yki target, *ex-lacZ* (βGal) reporter and Yki levels. (C) Example of a *wts^RNAi^* mosaic disc with RNAi-expressing clones being GFP-positive. *wts* knockdown leads to upregulation of the Yki target, *ex-lacZ* (βGal) reporter and Yki levels. (D) Example of a *Vha44* overexpression mosaic disc, with Vha44 overexpressing clones being GFP-positive. *Vha44* overexpression leads to upregulation of the Yki target, *ex-lacZ* (βGal) reporter and Yki levels. (A’, B’, C’, D’) Higher magnification examples of *ex-lacZ* (βGal) stainings in GFP-positive clones for the different samples. (E) Quantification of the *ex-lacZ* (βGal) pixel intensity ratio of the transgenic clones compared to *wild-type* clones. Error bars represent SEM. **** P-value<0.0001 *** P-value=0.0009 (one-way ANOVA with Bonferroni post-test). The scale bar represents 50 μm and in Á-D’ the scale bar represents 10 μm. Posterior is to the left in all images.

### Vap33 activates the Hippo pathway

Since we have previously shown that Lgl functions with Vap33 in negatively regulating the V-ATPase [25], we then tested if Vap33 is also involved in Hippo pathway regulation. As we have previously reported [5], in *lgl* mutant clones the levels of the Hippo pathway target, Diap1, are increased (**Fig 2A**, quantified in **Fig 2G**), and a distorted adult eye phenotype is observed (**Fig 2B**). We found that when Vap33 was overexpressed in clones, the expression of Diap1 was reduced (**Fig 2C**, quantified in **Fig 2G**), and adult eyes appeared only slightly disorganised (**Fig 2D**). However, overexpression of *Vap33* in *lgl* mutant clones resulted in a reduction of the elevated Diap1 protein levels (**Fig 2E**) that are observed in *lgl* mutant clones (**Fig 2A**) to below *wild-type* levels, similar to *Vap33* overexpression alone (**Fig C**, quantified in **Fig 2G**), and rescued the *lgl* mutant mosaic adult eye phenotype (**Fig 2F**), indicating that Vap33 is epistatic to Lgl. Together these results show that Vap33 activates the Hippo pathway and that Vap33 is epistatic to *lgl* impairment in the activation of the Hippo pathway.

**Fig. 2:**
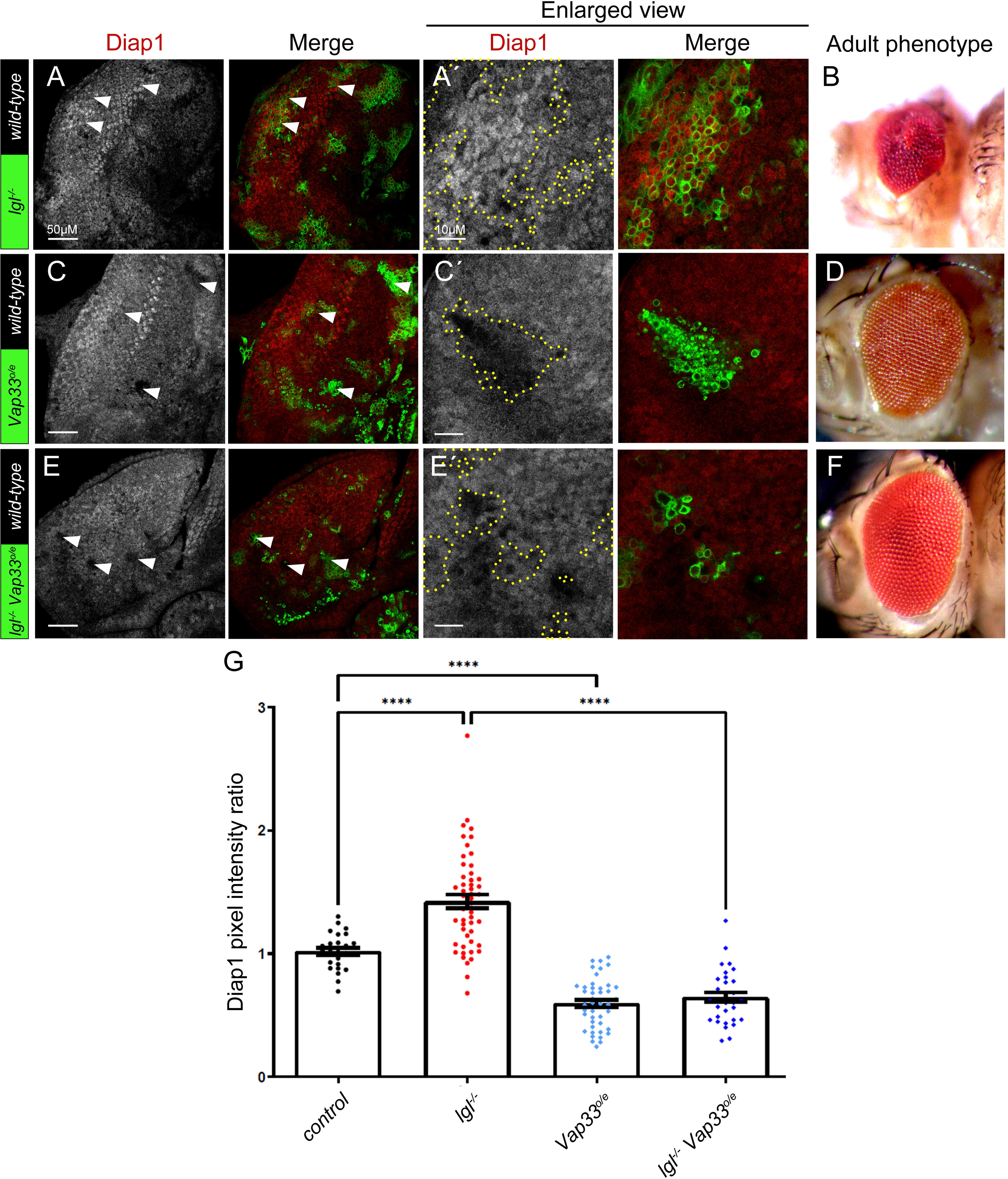
Vap33 activates the Hippo pathway. (A, C, E) Confocal planar sections of mosaic eye discs, stained for the Yki target Diap1 (grey, or red in merges). Mutant clones are GFP-positive, and examples are indicated by arrowheads. (A) Example of a *lgl*^-^ mosaic disc showing elevated Diap1 levels in the mutant clones. (B) *lgl*^-^ mosaic adult female eye. (C) Example of *Vap33^o/e^* mosaic disc showing reduced Diap1 levels. (D) *Vap33^o/e^*mosaic adult female eye. (E) Example of *lgl*^-^ *Vap33^o/e^*mosaic disc showing normalized Diap1 levels. (F) *lgl*^-^ *Vap33^o/e^*mosaic adult female eye. (Á, Ć, É) higher magnifications of Diap1 stainings for the different mosaic tissues. (G) Quantification of Diap1 pixel intensity ratio of mutant/transgenic clones compared with *wild-type* clones. Error bars represent SEM. **** P-value<0.0001 (one-way ANOVA with Bonferroni post-test). In A, C, E, the scale bar represents 50 μm, and in Á, Ć, É the scale bar represents 10 μm. Posterior is to the left in all images.

### Vap33 and Lgl form a protein interaction network with RtGEF (Pix), Git, Arf79F and Hpo

Since our results implicated Vap33 in the regulation of the Hippo pathway, we sought to determine whether this occurs via protein-protein interactions. Interestingly, the global human proteomics analysis has elucidated the protein interaction network of the human Vap33 orthologs, VAPA/B, and revealed amongst the interacting proteins the Hpo orthologs, MST1/2 (STK3/4) [29]. To determine whether *Drosophila* Vap33 also interacts with Hpo we conducted affinity purification-mass spectrometry (AP-MS) on endogenously expressed Vap33-YFP *in vivo* using immunoprecipitation with a GFP-nanobody [31]. We decided to use an *in vivo* approach since the common *Drosophila* cell line (S2 cells, thought to be derived from hemocytes) used for protein interaction analysis are not polarised cells, and by using *Drosophila* tissues, we hoped to reveal physiologically relevant interactors important in polarised epithelial tissues. We prepared proteins from Vap33-YFP-expressing embryos and conducted the experiment in triplicate together with controls. As expected, amongst the AP-MS Vap33 interactors we observed the V-ATPase component, Vha68-2 as a medium confidence interactor (significance analysis of interactome, SAINT score ∼0.77) and three other V-ATPase components as lower confidence interactors (**Supp File 1**). Surprisingly, Lgl and Hpo were not detected, suggesting that under the conditions used in embryonic cells these proteins do not form strong interactions with Vap33. However, pertinently, with respect to the Hippo pathway, we detected as high confident interactors (SAINT scores ∼1), the Hpo-interacting proteins, RtGEF (Pix, a Rho-type guanine nucleotide exchange factor) and Git (G protein-coupled receptor kinase interacting ArfGAP), are actin cytoskeletal regulators that form a protein complex [32, 33], and also bind to and activate Hpo [34].

To confirm these interactions, we conducted co-immunoprecipitation (co-IP) analyses in S2 cells. We co-transfected Vap33 tagged with V5 and RtGEF tagged with HA and conducted IP-western blot analysis (**Fig 3**). Vap33-V5 co-IPed with RtGEF-HA in both directions (**Fig 3A, B**). We also investigated whether Vap33 and Hpo could form a complex by co-transfecting Vap33-V5 and Hpo-Flag in S2 cells and conducting IP-Western blots. Indeed, Vap33 co-IPed with Hpo in both directions (**Fig 3C, D**). Thus, Vap33 binds to both RtGEF and Hpo, supporting our AP-MS data, and consistent with previous studies showing that Hpo interacts with RtGEF/Git [34], and that the human Vap33 orthologs, VAPA/B, interact with the Hpo orthologs STK3/4 [29].

**Fig 3:**
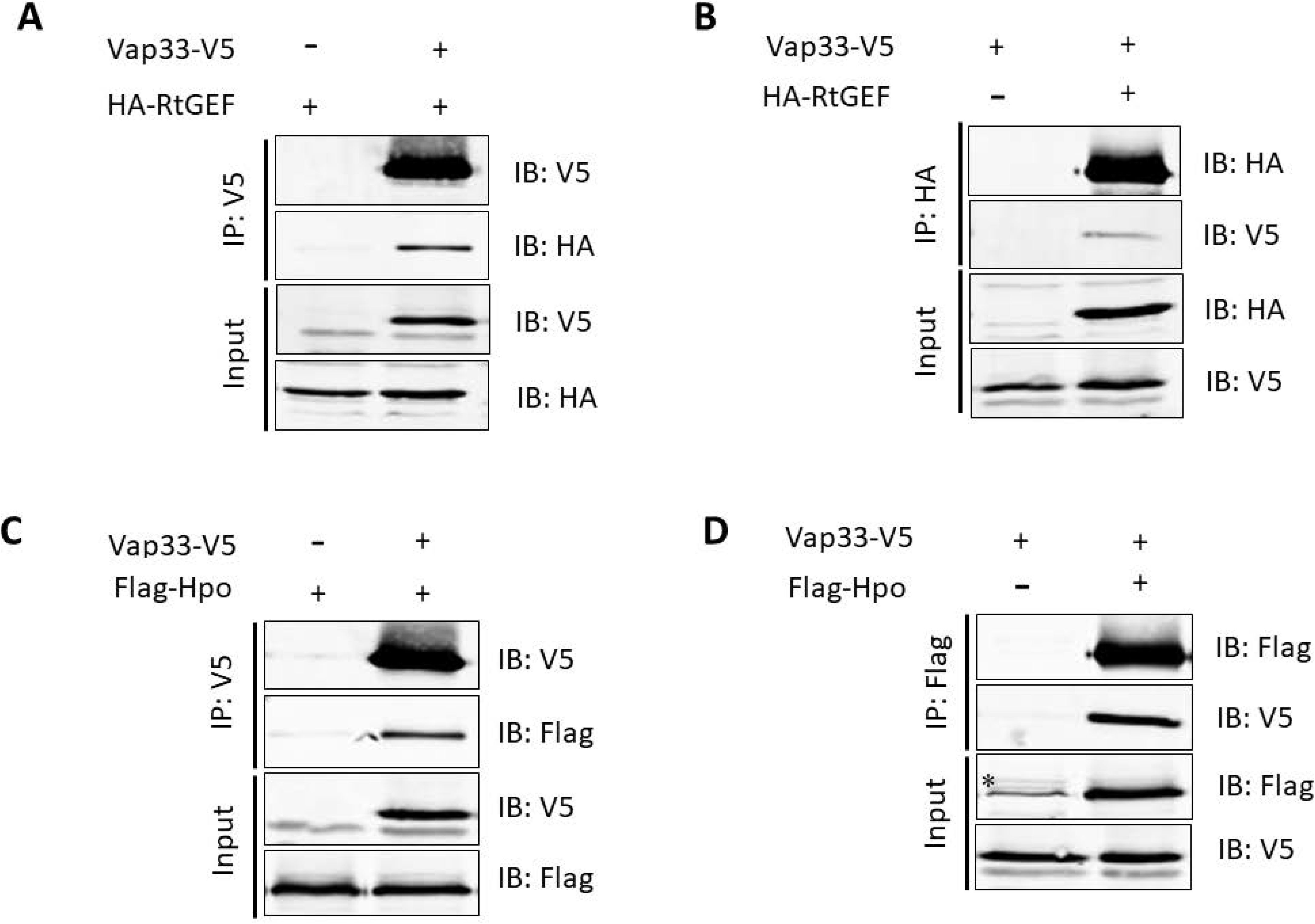
Vap33 interacts with RtGEF and Hpo in S2 cells. (A-D) Indicated protein constructs were expressed in S2 cells, and protein interactions were analysed by co-immunoprecipitation (co-IP). IP: immunoprecipitation antibody; IB: immunoblot antibody. Asterisk in (D) indicates non-specific band.

We then examined whether interactions between Lgl or Vap33 and RtGEF/Git occurred in *Drosophila* larval epithelial tissues, by using the proximity ligation assay (PLA) [27]. First, using larval tissues (eye-antennal epithelial, salivary gland and brain) from endogenously GFP-tagged Lgl flies [25, 35] and antibodies against GFP and Vap33, we confirmed that PLA foci were observed for Lgl and Vap33 (**Fig 4A, Supp Fig 1G, 1H**) [25], suggesting that these proteins physically interact *in vivo*. We also observed foci for Lgl-GFP and atypical protein kinase C (aPKC), by using GFP and aPKC antibodies in eye-antennal discs and salivary gland cells (**Supp Fig 1A, C**). Importantly, no PLA signals were observed in the single antibody negative controls (**Supp Fig 1B, 1D-F**). We then conducted PLAs on eye-antennal epithelial and salivary gland tissues using flies expressing the endogenously tagged Lgl-GFP and Git-RFP transgene [36], using GFP and RFP antibodies, which revealed multiple PLA foci (**Fig 4B, Supp Fig 1J**). Likewise, PLAs performed on Git-RFP transgenic fly eye-antennal epithelial and salivary gland tissues using RFP and Vap33 antibodies, revealed multiple PLA foci (**Fig 4C, Supp Fig 1K**). Thus, PLA data confirmed that Git interacts *in vivo* with Vap33 and Lgl in the *Drosophila* larval eye-antennal epithelium (**Fig 4**), as well as other polarised cells in the salivary gland and brain (**Supp Fig 1**).

**Fig. 4:**
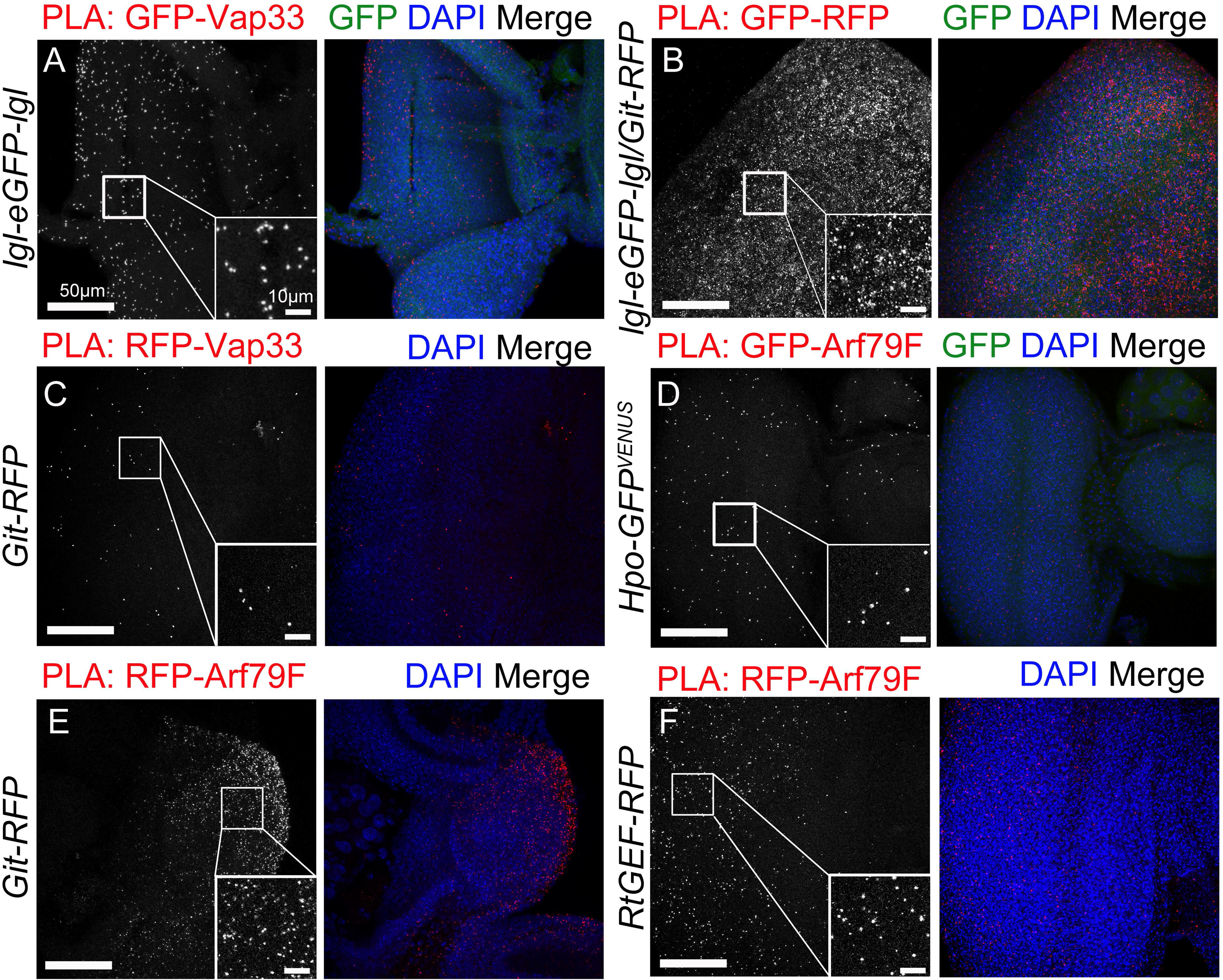
Vap33 interacts with Git, Arf79F interacts with Git, RtGEF and Hpo, and Lgl interacts with Vap33 and Git in vivo. (A-F) Confocal planar images showing *in situ* proximity ligation assays (PLAs) in third instar larval eye discs. Positive PLA results between the indicated proteins are visualized by punctate signal (grey or red in the merges). Nuclei are stained with DAPI (blue). Insets show high magnification images of the PLA foci. (A) Positive-control PLA in *lgl-eGFP-lgl* eye discs using antibodies against GFP and Vap33. (B) PLA in *lgl-eGFP-lgl*/*Git-RFP* eye discs using antibodies against RFP and GFP. (C) PLA in *Git-RFP* eye discs using antibodies against RFP and Vap33. (D) PLA in *Hpo-GFP* eye discs using antibodies against GFP and Arf79F. (E) PLA in *Git-RFP* eye discs using antibodies against RFP and Arf79F. (F) PLA in *RtGEF-RFP* eye discs using antibodies against RFP and Arf79F. Scale bars represent 50 μm.

Interestingly, in mammalian cells, Git interacts with the ADP ribosylation factor, Arf1 and inhibits its activity [33], and examination of the *Drosophila* Hippo pathway proteomics dataset [37], revealed that the *Drosophila* Arf1 ortholog, Arf79F, is a high confidence interactor with Hpo (SAINT score >0.8), which was validated by co-IP analysis [37]. Based on these physical interactions, we also tested by PLA whether Arf79F interacted with Git, RtGEF and Hpo. Indeed, using flies containing a Hpo-GFP-Venus transgene [38] and antibodies against Arf79F, we confirmed via PLA that Hpo and Arf79F interact in eye-antennal epithelial and brain tissue (**Fig 4D, Supp Fig 1I**). Similarly, using PLAs on eye-antennal discs and salivary glands from Git-RFP or RtGEF-RFP transgenic flies [36] and RFP and Arf79F antibodies, Arf79F was confirmed to interact with Git and RtGEF **(Fig 4E, 4F, Supp Fig 1L, M**). Thus, we have revealed that Arf79F binds to Git, RtGEF and Hpo in eye-antennal tissue (**Fig 4**), as well as salivary gland or brain tissues (**Supp Fig 1**).

### Vap33 overexpression rescues increased Hippo pathway target gene expression in *RtGEF* mutant tissue

To determine if *Vap33* genetically interacts with *RtGEF*, we examined the expression of the Hippo pathway target, Diap1, in *RtGEF* mutant (*RtGEF^P1036^*) clones that overexpress Vap33. Individually, *RtGEF* mutant clones showed increased Diap1 expression (**Fig 5A**), as expected [34], and resulted in a slightly enlarged adult eye phenotype (**Fig 5B**). The elevated Diap1 expression observed in *RtGEF* mutant clones was rescued to normal levels upon *Vap33* overexpression (**Fig 5C**, quantified in **Fig 5E**), and the adult eye size was normalized (**Fig 5D**). Thus, consistent with the protein interactions, RtGEF and Vap33 genetically interact in the regulation of the Hippo pathway in a manner consistent with both Vap33 and RtGEF acting to induce Hippo pathway activity.

**Fig. 5:**
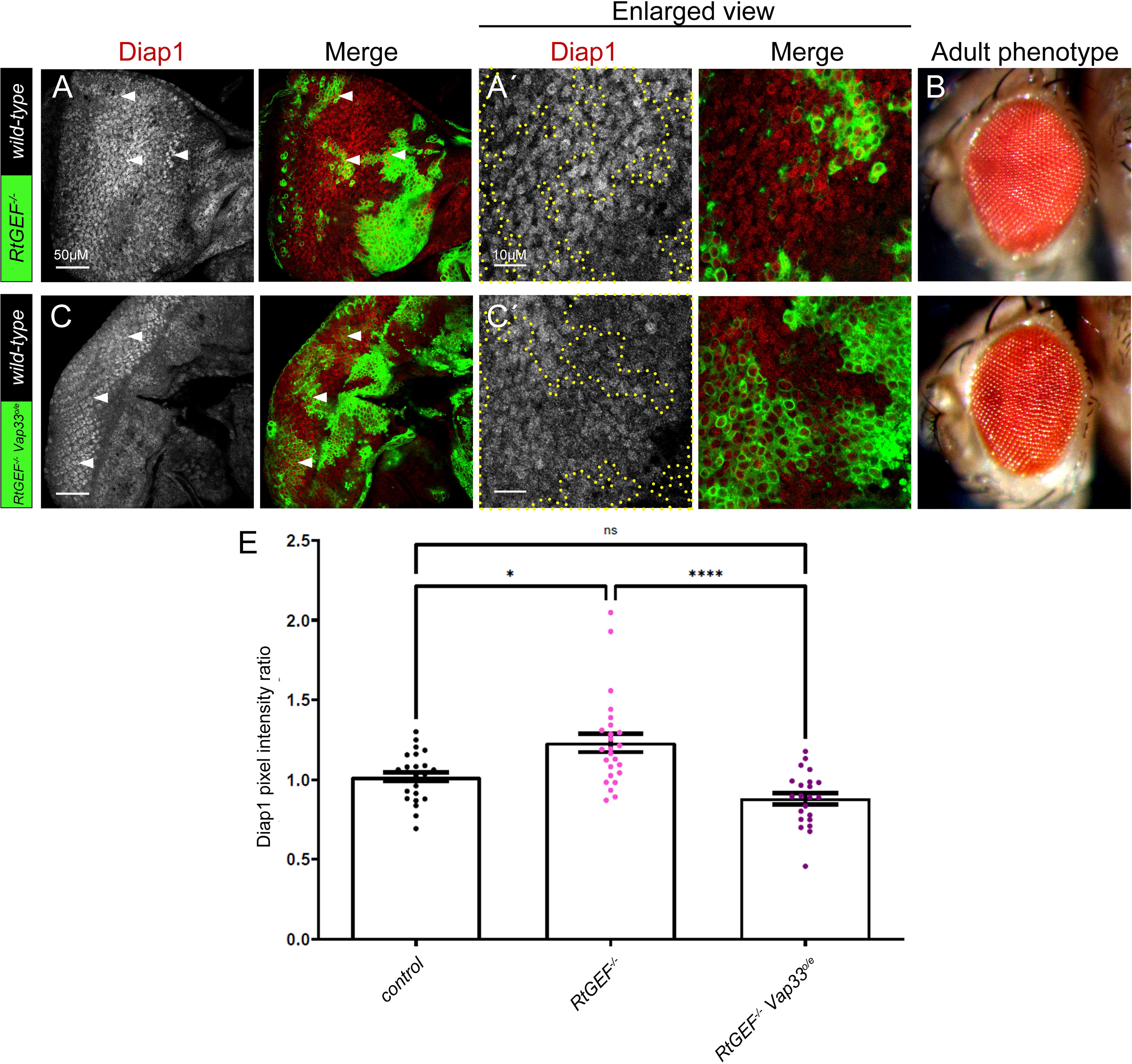
Vap33 overexpression rescues increased Hippo pathway target gene expression in *RtGEF* mutant tissue. (A) Confocal planar section of a *RtGEF^-^* mosaic disc stained for the Yki target Diap1 (grey, or red in merges, mutant clones are GFP-positive, examples indicated by arrowheads). (B) *RtGEF^-^* mosaic adult female eye. (C) Confocal planar section of *RtGEF^-^ Vap33^o/e^*mosaic disc stained for Diap1 (grey, or red in merge, mutant tissue is GFP-positive, examples indicated by arrowheads). (D) *RtGEF^-^ Vap33^o/e^* mosaic adult female eye. (Á, Ć) higher magnifications of Diap1 stainings of the respective samples. (E) Quantification of Diap1 pixel intensity ratio of mutant/transgenic clones compared to *wild-type* clones. Error bars represent SEM. **** P-value<0.0001 * P-value=0.019 n.s., differences not significant (one-way ANOVA with Bonferroni post-test). Posterior is to the left. Scale bar represents 50 μm (A, C) and 10 μm (A’ C’).

### Git knockdown rescues the reduced Hippo pathway target gene expression in *Vha68-2* mutant clones

Next, we examined genetic interactions between Git and a component of the V-ATPase, Vha68-2, in Hippo pathway regulation. As we have previously observed [25], *Vha68-2* mutant clones were small, and they showed reduced expression of the Hippo pathway target, Diap1 (**Fig 6C**) relative to *wild-type* clones (**Fig 6A**, quantified in **Fig 6G**), and resulted in reduced and disorganised adult eyes relative to the *wild-type* control (**Fig 6D** compared with **Fig 6B**). When Git was knocked down using a transgenic RNAi line in *Vha68-2* mutant clones, partial rescue of the reduced Diap1 expression was observed (**Fig 6E**, quantified in **Fig 6G**) and the disorganised *Vha68-2* mutant mosaic adult eye phenotype was strongly rescued (**Fig 6F**). These results suggest that the activation of Hippo signalling (reduced Diap1 expression) that occurs upon reducing V-ATPase activity depends upon Git.

**Fig. 6:**
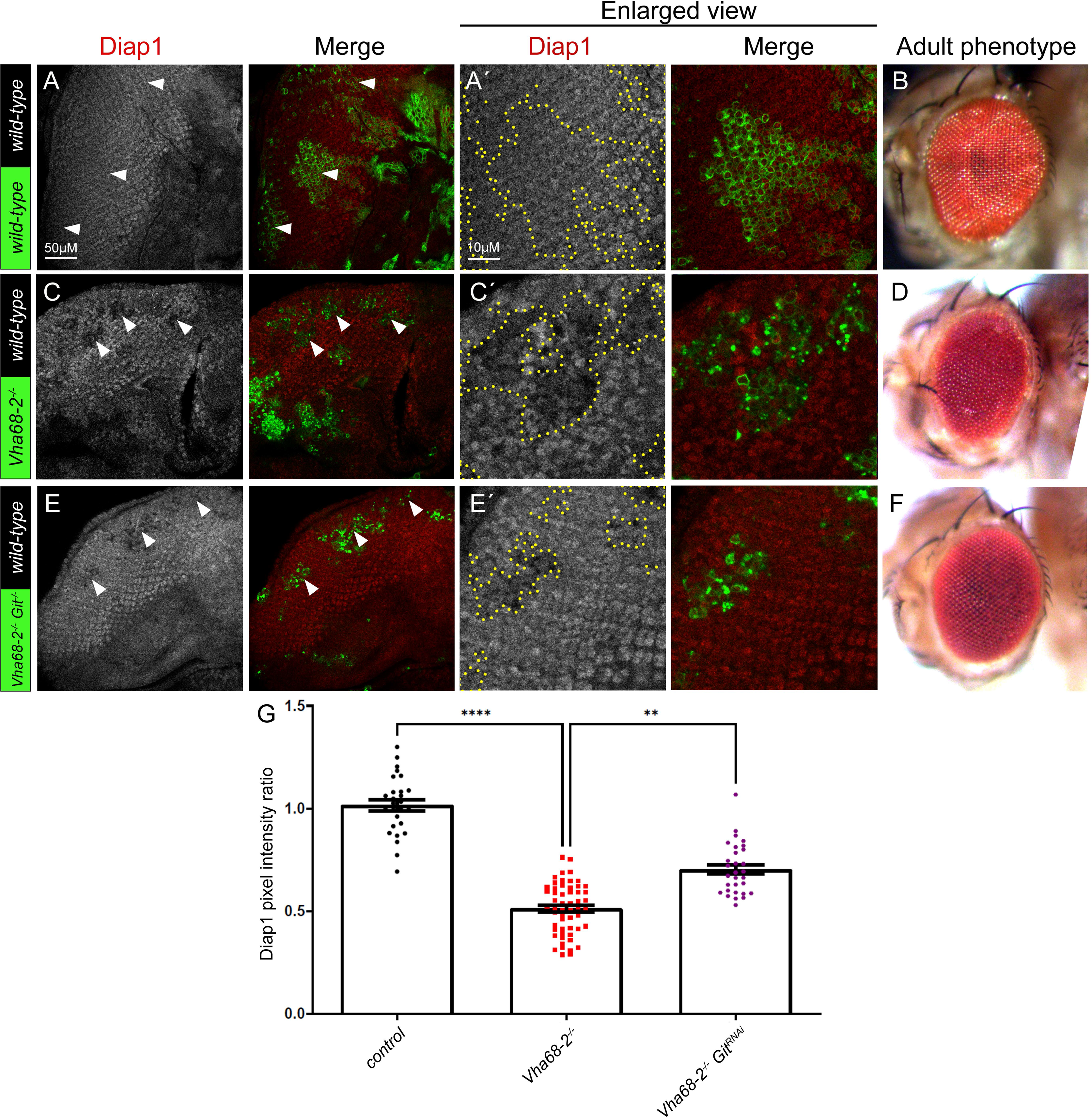
G*i*t knockdown rescues the reduced Hippo pathway target gene expression in *Vha68-2* mutant clones. (A) Confocal planar section of control mosaic eye discs (clones marked with GFP) stained for the Yki target Diap1 (grey, or red in merges, example GFP-positive clones indicated by arrowheads) showing endogenous expression of Diap1. (B) Control mosaic adult female eye. (C) Confocal planar section of *Vha68-2^-^‘*^-^ mosaic disc stained for Diap1 (grey, or red in merges, mutant clones are GFP-positive, examples indicated by arrowheads). (D) *Vha68-2^-^‘*^-^ mosaic adult female eye. (E) Confocal planar section of *Vha68-2^-^‘*^-^ *Git^RNAi^* mosaic disc stained for Diap1 (grey, or red in merge, mutant tissue is GFP-positive, examples indicated by arrowheads). (F) *Vha68-2 Git^RNAi^* mosaic adult female eye. (Á, Ć, É) Higher magnifications of Diap1 stainings of the corresponding samples. (G) Quantification of Diap1 pixel intensity ratio of the mutant/transgenic clones relative to *wild-type* clones. Error bars represent SEM. **** P-value<0.0001 ** P-value=0.054 (one-way ANOVA with Bonferroni post-test). In all images, posterior is to the left. Scale bar represent 50 μm (A, C, E), and 10 μm (Á, Ć, É).

### Arf79F is required for Lgl-mediated regulation of the Hippo pathway

We then tested whether Arf79F is required for the regulation of the Hippo pathway by Lgl. As we previously observed, *lgl* mutant tissue shows impaired Hippo pathway signalling, as revealed by the elevated levels of Diap1 (**Fig 7C** compared to the control, **Fig 7A**, quantified in **Fig 7M**) and results in a distorted, disorganised adult eye phenotype relative to the *wild-type* control (**Fig 7D** compared with **Fig 7B**). We therefore sought to determine whether *Arf79F* genetically interacts with *lgl* in its regulation of the Hippo pathway. When *Arf79F* was knocked down, using a RNAi line (which we showed effectively reduced Arf79F protein levels in eye disc clones, **Supp Fig 2A, B**), a decrease in Diap1 expression was observed and clones were smaller than *wild-type* clones **(Fig 7E**), suggesting that the *Arf79F* knockdown clones were being out-competed, consistent with Hippo pathway upregulation. However, the *Arf79F* adult mosaic eyes showed only slight disorganisation (**Fig 7F**). Knockdown of *Arf79F* in *lgl* mutant clones also resulted in low Diap1 levels, similar to *Arf79F* knockdown alone (**Fig 7G** compared with **Fig 7E**, quantified in **Fig 7M**), suggesting that *Arf79F* is epistatic to *lgl* in the regulation of the Hippo pathway. Consistent with this, the *lgl* mutant mosaic adult eye phenotype was normalised upon *Arf79F* knockdown (**Fig 7H**). Additionally, we used a dominant-negative version of the activator of Arf79F, Sec71 (*Sec71^DN^*), to reduce Arf79F function. When expressed individually in clones, *Sec71^DN^*had no significant effect upon Diap1 levels (**Fig 7I**, quantified in **Fig 7M)**, and did not obviously affect the adult eye phenotype (**Fig 7J**). However, when *Sec71^DN^* was expressed in *lgl* mutant clones, lower Diap1 levels were observed (**Fig 7K**, quantified in **Fig 7M),** and the *lgl* mutant mosaic adult eye phenotype was normalised (**Fig 7L**). Interestingly, although *Sec71^DN^* expression did not have a significant effect on Diap1 expression on its own, when expressed in *lgl* mutant clones it reduced Diap1 levels to below that of the control (**Fig 7M**), suggesting that Lgl depletion depends on Sec71 (and thus Arf79F) to inhibit Hippo signalling. These results show that, consistent with the binding of Arf79F with Hpo (**Fig 4D**), knockdown of Arf79F reduces Diap1 expression, suggesting that Arf79F acts to inhibit the Hippo pathway. Furthermore, knockdown of Arf79F (or expression of *Sec71^DN^*) rescued the elevated expression of Diap1 in *lgl* mutant clones (**Fig 7M),** indicating that Arf79F levels/activity are required for the inhibition of Hippo signalling in *lgl* mutant tissue. Thus, Arf79F is a negative regulator of Hippo signalling that acts downstream of Lgl. Since mammalian Git proteins act to inhibit Arf1 activity [33], our results are consistent with a mechanism where Lgl and Vap33 promote RtGEF/Git activity to inhibit Arf79F (Arf1), and thereby activate the Hippo pathway.

**Fig. 7:**
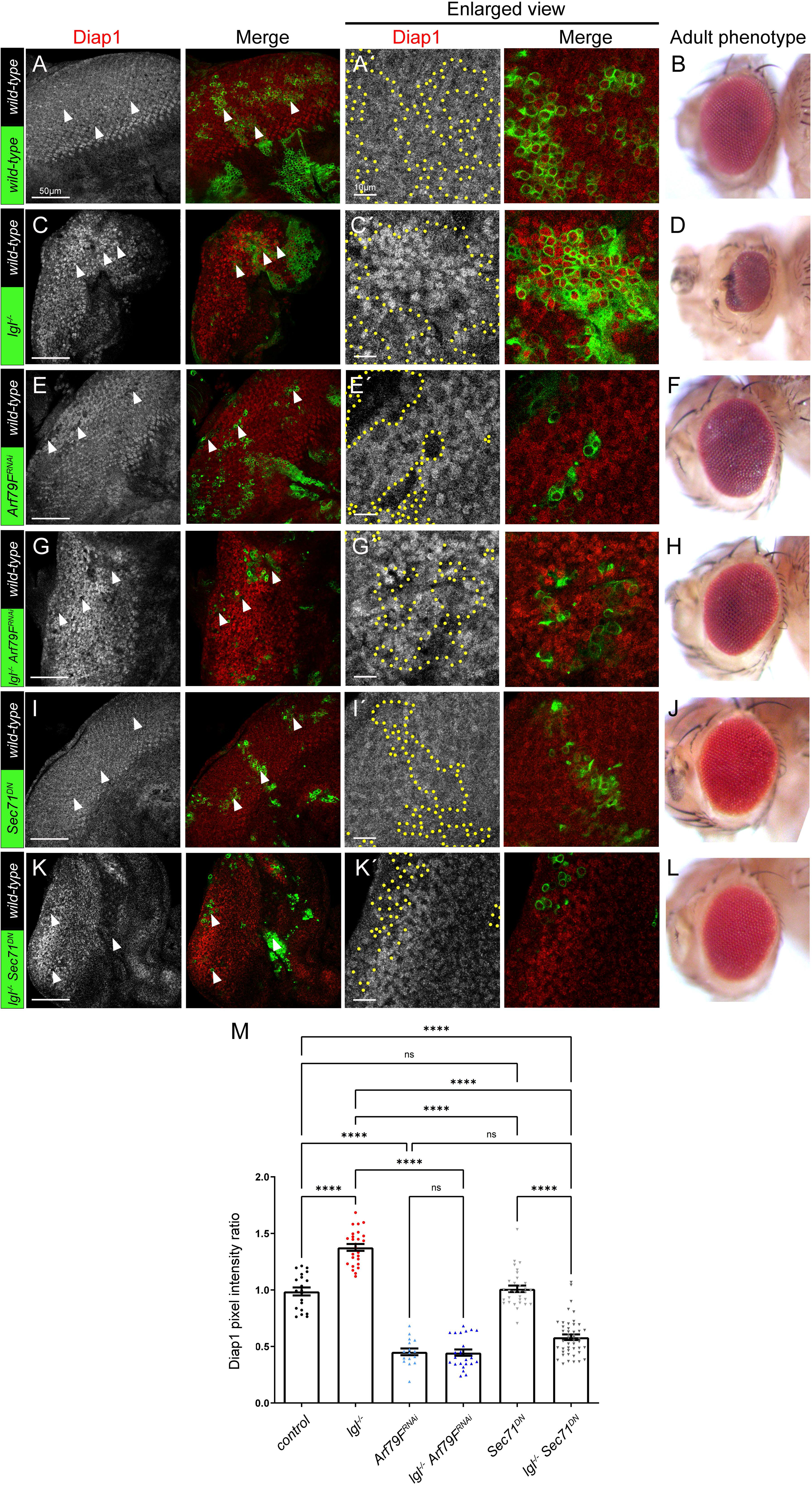
Knockdown of Arf79F prevents the upregulation of Hippo pathway targets. (A, C, E, G, I, K) Confocal planar images of mosaic third instar larval eye-antennal discs, stained for the Yki target, Diap1 (grey, or red in merge). In all instances, example GFP-positive clones are marked by arrowheads. (A) Control eye discs (clones marked by GFP) showing endogenous expression of Diap1. (B) Control mosaic adult female eye. (C) *lgl*^-^ mosaic disc showing elevated Diap1 expression in mutant clones (GFP-positive). (D) *lgl*^-^ mosaic adult female eye. (E) *Arf79F^RNAi^* mosaic disc showing reduced Diap1 expression in transgene-expressing clones (GFP-positive). (F) *Arf79F^RNAi^* mosaic adult female eye. (G) *lgl*^-^ *Arf79F^RNAi^* mosaic disc showing reduced Diap1 expression in mutant/transgene-expressing clones (GFP-positive). (H) *lgl*^-^ *Arf79F^RNAi^* mosaic adult female eye. (I) *Sec71^DN^* mosaic disc showing normal Diap1 expression in transgene-expressing clones (GFP-positive). (J) *Sec71^DN^* mosaic adult female eye. (K) *lgl*^-^ *Sec71^DN^* mosaic disc showing reduced Diap1 expression in mutant/transgene-expressing clones (GFP-positive). (L) *lgl*^-^ *Sec71^DN^* mosaic adult female eye. (Á, Ć, É, G’, Í, K’) higher magnifications of Diap1 stainings in the corresponding samples. (M) Quantification of Diap1 pixel intensity ratio of mutant/transgene clones relative to *wild-type* clones. Error bars represent SEM. **** P-value<0.0001 (one-way ANOVA with Bonferroni post-test). Posterior is to the left in all images. Scale bars represents 50 μm (A, C, E, G, I, K), or 10 μm (Á, Ć, É, G’, Í, K’).

## Discussion

In this study we have revealed a new mechanism for the control of the Hippo pathway by the cell polarity regulator, Lgl. We show herein that V-ATPase activity leads to inhibition of Hippo signalling, and conversely that Vap33 activates the Hippo pathway downstream of Lgl. We discovered that Vap33 physically interacts with RtGEF and Git using *in vivo* affinity-purification mass spectrometry. RtGEF and Git are Hpo interactors [36], and we confirmed that Vap33 interacts with RtGEF and Hpo in S2 cells, and that Vap33 and Lgl are in close proximity with Git in *Drosophila* cells. We also show that the ADP ribosylation factor, Arf79F, the mammalian ortholog of which (ARF1) binds to the mammalian Git orthologs, GIT1/2 [33], and is in close proximity with Git, RtGEF and Hpo in *Drosophila* cells. Functionally, we show that overexpression of Vap33 rescues the elevated Diap1 levels (impaired Hippo signalling) in *RtGEF* mutant clones. Furthermore, the reduced Diap1 expression (elevated Hippo signalling) upon V-ATPase impairment (*Vha68-2* mutant clones) is rescued by knockdown of Git. We also show that Arf79F knockdown leads to reduced Diap1 expression (increased Hippo signalling) downstream of Lgl. Altogether, our data are consistent with a model where Lgl and Vap33 activate the Hippo pathway by a dual mechanism: 1) Lgl and Vap33, through interaction with RtGEF/Git/Arf79F, positively regulate Hippo pathway activity, and 2) Lgl and Vap33, through interaction with components of the V-ATPase, block V-ATPase activation and prevent its negative regulation of the Hippo pathway **(Fig 8)**. Precisely how the V-ATPase functions to regulate Hippo signalling is unknown. However, the V-ATPase acts to increasing vesicle acidification, and this has been shown to impact the efficacy of many signalling pathways [39-41]. The V-ATPase might therefore act to inhibit Hippo pathway signalling by blocking the interaction of Lgl/Vap33/RtGEF/Git/Arf79F with Hpo in vesicles, thereby altering Hpo localization and inhibiting its activity. Consistent with this notion, we observed that Lgl colocalizes with endocytic vesicle markers [18], and that Hpo localization is altered in *lgl* mutant tissue [5]. Furthermore, the subcellular localization of Hpo has been shown to be important for its activation and for Hpo-mediated phosphorylation and activation of its downstream protein kinase, Wts [42].

**Fig. 8:**
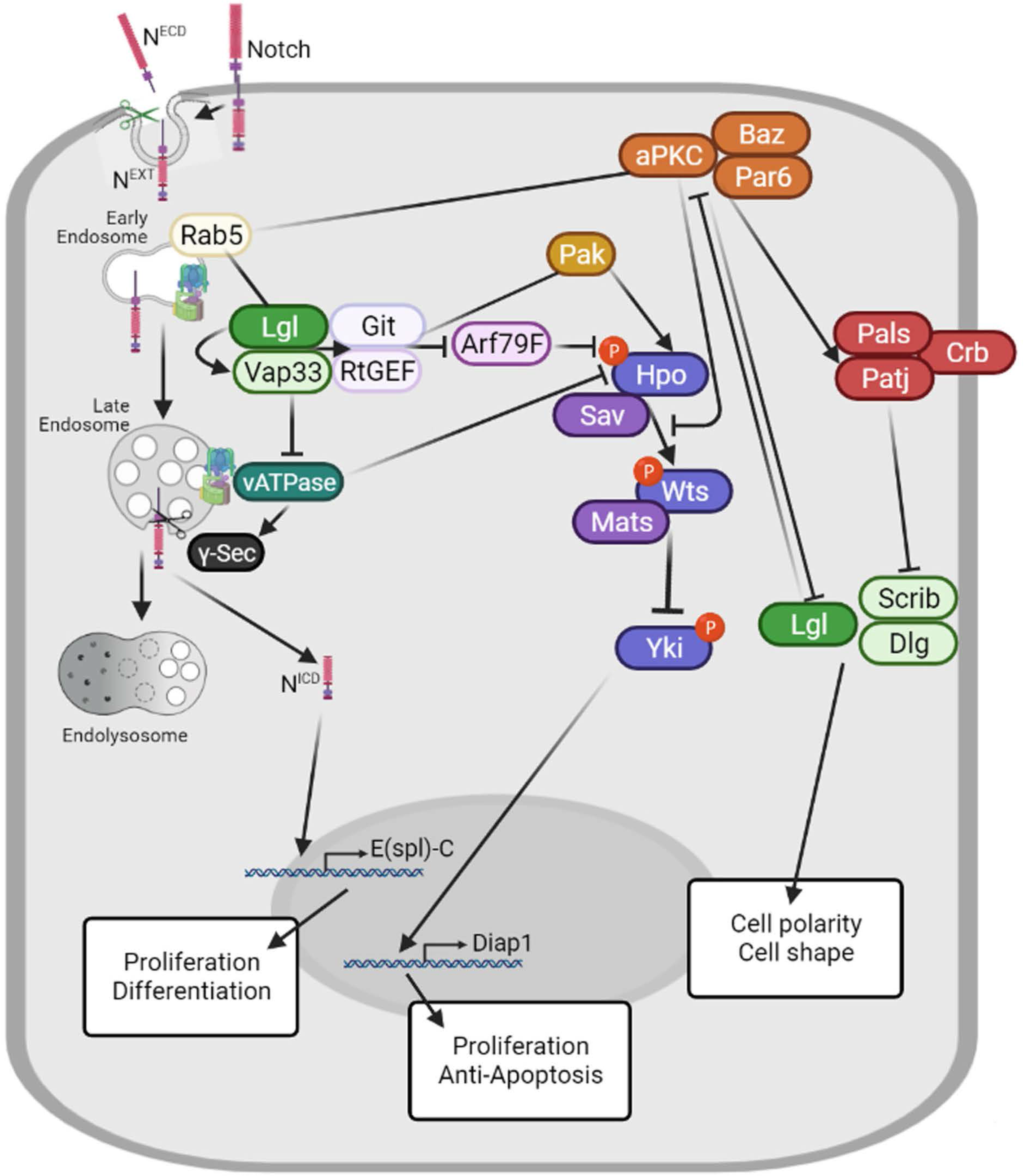
Model for the regulation of Hippo and Notch signalling by Lgl/aPKC. Lgl/Vap33 have a dual action in activating Hippo signalling: 1) through binding to RtGEF/Git, which opposes Arf79F inhibition of Hpo, and 2) by binding to V-ATPase components and inhibiting V-ATPase activity, which is an inhibitor of Hippo signalling. The Lgl interactor, Rab5, may be involved to link Lgl and Vap33 to early endosomes. It is possible that the RtGEF/Git interacting protein kinase, Pak (Pak1, Pak3, Mbt), is involved in activating Hpo downstream of Git/RtGEF. By reducing V-ATPase activity, Lgl/Vap33 inhibits Notch signalling by decreasing the production of the active N^ICD^ isoform by γ-Secretase in endosomes, thereby leading to lower expression of Notch targets, such as *E(Spl)-C* genes. Lgl also opposes aPKC activity, and in mammalian cells elevated PKC inhibits Hippo signalling by sequestering Hpo (MST1/2) away from its target, Wts (LATS).

In their interactions with the Hippo pathway, Git and RtGEF function to activate Hpo [34], while our analyses indicate that Arf79F acts to inhibit Hippo signalling, since elevated Hippo pathway activity (increased Diap1 expression) in *lgl* mutant clones was normalised upon knockdown or inhibition of Arf79F. Previous studies examining the interactions between mammalian orthologues of Git and Arf79F, showed that GIT1/2 acts to inactivate ARF1 [33], and therefore it is likely that Git (and RtGEF) act to block Arf79F activity to mediate Hpo activation (**Fig 8**). Lgl and Vap33 might then function to promote Git/RtGEF inhibition of Arf79F, facilitating activation of Hpo. Arf79F (ARF1) is a regulator of vesicular trafficking [43], and may inhibit Hpo by relocalizing it away from its activators, such as Expanded and Fat, at specific apical subcellular domains [42]. Interestingly, from our previous Lgl interactome analysis, Arf79F can be linked to Lgl through the Arf79F-binding protein Rab5 [28, 37], which also binds to Lgl and aPKC at high confidence, SAINT score ∼1) ([25] and unpublished data). Thus, it is possible that the early endosomal regulator, Rab5, may also be involved in linking Lgl/Vap33 and aPKC to Arf79F in early endosomes, where they may also interact with Vap33, Git/RtGEF and Hpo.

It is unclear whether the RtGEF/Git regulated protein kinase, Pak [33], is involved in the regulation of Hippo signalling, since although individual knockdown of two Pak paralogs in *Drosophila*, Pak1 and Pak3, did not affect Hippo pathway signalling [34], it is possible that there may be redundancy between Pak1, Pak3, and the third *Drosophila* Pak paralog, Mushroom bodies tiny (Mbt) [44-47], in the regulation of Hippo pathway signalling. Pak proteins have been shown to be involved in cell polarity, F-actin regulation and morphogenesis in *Drosophila* [45-54], however Mbt also plays a role in tissue growth during *Drosophila* developmental [44, 55], suggesting Mbt is a candidate for investigation of a potential role downstream of RtGEF/Git in Hippo pathway regulation.

Our analyses here, and in our previous study [25], have revealed that Lgl interacts with Vap33 to positively regulate the Hippo pathway and to negatively regulate the Notch pathway. Mechanistically, Lgl/Vap33 inhibits the Notch pathway by inhibiting V-ATPase activity and reducing vesicle acidification that is required for the cleavage of the Notch receptor and release of Notch^ICD^ from vesicles [25], where it can then translocate into the nucleus and promote transcription of its target genes (**Fig 8**). Interestingly, Arf79F (Arf1) is required for Notch signalling in *Drosophila* hemocyte differentiation, where it promotes Notch trafficking [56]. Thus, it is possible that Lgl/Vap33/RtGEF/Git may also inhibit Arf79F to reduce Notch signalling, and that in *lgl* mutant tissue elevated Arf79F activity may also contribute to the elevated Notch activation. Indeed, we found that Arf79F is required for the elevated Notch pathway signalling in *lgl* mutant clones, since *Arf79F* knockdown in *lgl* mutant clones normalized the expression of the Notch pathway reporter, *E(spl)m8-lacZ* (**Supp Fig 3**). Whether RtGEF/Git are also regulators of Notch signalling remains to be determined.

In summary, our study has revealed a new mechanism for Hippo pathway regulation by the cell polarity and tumour suppressor protein, Lgl. Mechanistically, Lgl and Vap33 function in a dual manner to promote Hippo signalling by both reducing V-ATPase activity, and thereby preventing V-ATPase from inhibiting Hippo signalling and by acting through RtGEF/Git to prevent Arf79F from inhibiting Hpo. By promoting Hippo signalling as well as inhibiting Notch signalling, Lgl/Vap33 function to limit tissue growth. Mutation of Lgl therefore results in tissue overgrowth due to inhibition of the Hippo pathway and activation of the Notch pathway. This mechanism may be important in limiting tissue growth during development and may play a role in tissue homeostasis, enabling cell proliferation to occur after tissue damage, thereby enabling wound repair. Indeed, Hippo pathway signalling is inhibited, and Notch pathway signalling is activated after tissue wounding and both pathways are involved in tissue regeneration [57-60]. However, the involvement of Lgl/Vap33/Git/RtGEF/Arf79F and the V-ATPase in the regulation of Hippo and Notch signalling during the response to tissue wounding remains to be determined. Furthermore, whether the mammalian orthologs of Lgl and Vap33 also act via these mechanisms to control tissue growth in mammals remains to be determined.

## Experimental Procedures

### *Drosophila* stocks and husbandry

Fly stocks were generated in house or obtained from other laboratories or stocks centers as detailed in **Table S1**. All *Drosophila* genotypes for all the samples analysed in the figures and detailed in **Table S2**. All fly stocks and crosses were raised and undertaken on a standard cornmeal/molasses/yeast medium within temperature-controlled incubators at 25°C.

### Clonal Analysis

Mosaic analysis with a repressible cell marker (MARCM (GFP+)) [61] clones were generated as previously described [5], using the following stock: *ey-FLP, UAS-GFP; Tub-GAL80, FRT40A; Tub-GAL4/TM6B* MARCM 2L. Crosses were set up and left overnight at room temperature. Adults were then turned daily into new vials and allowed to lay for ∼24 hours at 25°C. L3 animals were then collected after ∼144 hours (6 days after the egg laying period). Samples were collected and prepared as described across multiple days to be similarly aged, and then stored at 4°C in 80% glycerol as necessary prior to mounting.

Heat shock induced Flip-out clones were generated using the stock: *hs-FLP; ex-lacZ; Act>CD2>GAL4, UAS-GFP.* Specifically, clones were induced by heat shocking larvae at 37°C for 15 minutes roughly 72 hours after egg laying. L3 imaginal discs were then dissected and prepared as described 72h after clone induction.

### Immunofluorescence

Third-instar larval eye-antennal discs, were dissected in phosphate-buffered saline (PBS), fixed in 4% paraformaldehyde for 30min, washed in PBS + 0.1 or 0.3% Triton X-100 (PBT), and blocked in PBT + 1% BSA (PBT/BSA). The tissues were incubated with the primary antibody in PBT/BSA over night at 4°C. After washing off the primary antibodies the tissues were incubated with the secondary antibodies in PBT for 1h at room temperature. After washing off the secondary antibodies the samples were mounted in 80% glycerol or Vectashield (Vector Laboratories).

Antibodies used were; mouse β-galactosidase (Sigma, 1:500), rabbit GFP (Invitrogen A11122, 1:500), mouse GFP (Invitrogen A11120, 1:500); mouse RFP (Invitrogen RF5R, 1:100) rabbit Yki (gift from K. Irvine, 1:400), mouse Diap1 (gift from B. Hay, 1:100), rabbit Vap33 (gift from H. Bellen, 1:1000), rabbit aPKC (Santa Cruz, 1:1000), rabbit Arf79F (gift from M.S. Inamdar, 1:500 [56]). Guinea pig Arf79F (gift from F. Yu, 1:200 [62]).

Secondary antibodies used were; anti-mouse Alexa 568, 633, 647, anti-rabbit Alexa 568, 633, and anti-guinea pig Alexa 568, 633 (1:500). DNA was stained with 2-(4-amidinophenyl)-1H-indole-6-carboxamidine (DAPI, 1 µM).

### Affinity Purification-Mass Spectrometry

Flies expressing endogenously YFP-tagged Vap33 (line #115288, Kyoto Stock Center) were grown in population cages, with embryos laid overnight collected on apple juice/agar plates. Extraction and protein purification were carried out as described in Neumuller et al. [31]. Briefly, extracts were prepared using Default Lysis Buffer (DLB) (50 mM Tris pH 7.5, 125 mM NaCl, 5% glycerol, 0.2% IGEPAL, 1.5 mM MgCl_2_, 1 mM DTT, 25 mM NaF, 1mM Na_3_VO_4_, 1mM EDTA, and 2x Complete protease inhibitor, Roche), spun down to clarify, incubated first with empty agarose beads and then with GFP-Trap resin (ChromoTek), followed by washes and elution in SDS sample buffer. The eluates were loaded on SDS-polyacrylamide gel and electrophoresed so that the dye front migrated ∼1 cm in the separating gel. The gel was then stained with Coomassie blue, and the lane was cut into two 5 mm x 5 mm pieces and sent for mass spectrometry analysis (Taplin Mass Spectrometry Facility, Harvard Medical School). Samples were digested with trypsin in-gel and peptides were analyzed using a Thermo Scientific Orbitrap mass spectrometer. The mass spectrometry analysis was conducted on three biological replicates for both experimental samples (*Vap33-YFP*) and controls (*yw* fly line). Data were analysed using the Significance Analysis of Interactome (SAINT) program [63], and a complete table of Vap33-YFP-interacting proteins is included as **Supplemental File 1**. Mass spectrometry data were deposited to the ProteomeXchange Consortium via the PRIDE partner repository [64] with the data set identifier PXD035110.

### Co-immunoprecipitation (co-IP) and Western Blot Analysis

Constructs used were: pMT-Vap33-V5 [25], pAc5.1-Flag-Hpo [34] and pAC5-1-HA-RtGEF [34]. *Drosophila* S2 cells were maintained in standard Schneider’s S2 medium with fetal bovine serum (Gibco) at 25°C, and transfections were performed using Effectene transfection reagent (Qiagen). CuSO_4_ was added to culture media at a final concentration of 0.35 mM for inducing expression of Vap33-HA. Cells were lysed using DLB (see above) and spun down to remove debris. Clear cell lysates were incubated with anti-V5, anti-HA, or anti-Flag beads (Sigma) for 2 hrs at 4°C. Beads were washed three times with lysis buffer, and protein complexes were eluted with SDS buffer. Proteins were separated via SDS-PAGE, blotted, incubated with primary and secondary antibodies, and signal was detected using the Odyssey imaging system (LI-COR). Primary antibodies used were: mouse anti-V5 (Sigma, 1:1,000), rabbit anti-Flag (Sigma, 1:1,000), rabbit anti-HA (Sigma, 1:1,000). Secondary antibodies used were: IRDye 800CW Donkey anti-Rabbit IgG (LI-COR), IRDye 680CW Donkey anti-Mouse IgG (LI-COR).

### Proximity ligation assay

The interactions between Lgl and aPKC, Vap33 or Git; Arf79F and Git, RtGEF or Hpo; and Vap33 and Git in *Drosophila* larval tissues were detected *in situ* using the Duolink® In Situ Red Starter Kit Mouse/Rabbit (Sigma, DUO92101) according to the instructions of the manufacturer. Briefly, primary antibody incubation was applied using the same conditions as immune-histofluorescence staining. Duolink secondary antibodies against the primary antibodies were then added. These secondary antibodies were provided as conjugates to oligonucleotides that were able to form a closed circle through base pairing and ligation using Duolink ligation solution when the antibodies were in close proximity [27] (a distance estimated to be <40 nm [65]). The detection of the signals was conducted by rolling circle amplification using DNA polymerase incorporating fluorescently labelled nucleotides into the amplification products. The resulting positive signals were visualized as bright fluorescent dots, with each dot representing one interaction event. As technical negative control one of the primary antibodies was not added therefore, no positive signals were obtained from that assay. An additional negative control was performed in a tissue without one of the antigens (GFP) and the full protocol was performed in those tissues. As a positive control, antibodies against two well-known interactors in the tissues, Lgl-aPKC and Lgl-Vap33, were used. The tissues were visualized using confocal microscopy (Zeiss Confocal LSM 780 PicoQuant FLIM or Zeiss LSM 800 Airyscan laser scanning confocal).

Primary antibody pairs used were mouse GFP with rabbit Vap33 (**Fig. 4A, S1G, H**), mouse RFP with rabbit GFP (**Fig. 4B, S1J**), mouse RFP with rabbit Vap33 (**Fig. 4C, S1K**), mouse GFP with rabbit Arf79F (**Fig. 4D, S1I**), mouse RFP with rabbit Arf79F (**Fig. 4E, F, S1L, M**), mouse GFP with rabbit aPKC (**Fig. S1A, C**). A GFP-tagged version of Lgl was used to detect interactions with Vap33, since the Lgl and Vap33 antibodies were both raised in rabbit. GFP-tagged Hpo was also used since the available Hpo antibody was raised in rats, and PLA antibodies designed for use with rat-raised antibodies were unavailable. RFP-tagged versions of RtGEF and Git were used for similar reasons.

### Imaging

Fluorescent-labelled samples were mounted in 80% glycerol or Vectashield (Vector Laboratories) and analysed by confocal microscopy (LEICA TCS SP5, Zeiss Confocal LSM 780 PicoQuant FLIM or Zeiss LSM 800 Airyscan laser scanning confocal). Images were processed using Leica LAS AF Lite and Fiji (Image J 1.50e). Images were assembled using Photoshop 21.2.3 (Adobe). Adult eyes were imaged on a dissecting microscope using a Scitec Infinity1 camera. Images were processed, analysed, and assembled using some combination of LAS AF Lite (Leica), Zen 2012 (Zeiss), Fiji and Photoshop 21.2.3 (Adobe).

### Statistical Analysis of Signal Intensity

Relative *Ex-lacZ* (**Fig. 1**), or *E(spl)m8-lacZ* (**Fig. S3**) βGal stainings, and Diap1 stainings (**Fig. 2, 5, 6, 7**)) within eye discs was determined using images taken at the same confocal settings. Average pixel intensity was measured using the measurement log tool from Fiji or Photoshop 5.1 (Adobe). In the case of *E(spl)m8-lacZ* analyses, clones were chosen just posterior to the morphogenetic furrow of each eye disc. Average pixel intensity was measured in mutant clones and the *wild-type* adjacent tissue of the same areas and expressed as a ratio of pixel intensity of the mutant clone relative to the *wild-type* tissue. To analyze and plot data, we used Microsoft Excel 2013 and GraphPad Prism 9. We performed a D’Agostino and Pearson normality test, and the data found to have a normal distribution were analysed by a two-tailed t test with Welch correction. In the case of multiple comparisons, we used a one-way ANOVA with Bonferroni post-test. Error bars represent SEM.

## Acknowledgments

We are grateful to all those who contributed fly stocks or antibodies to this study, to Bloomington, VDRC and NIG Stock Centers, OzDros and Flybase. HER was supported by a Senior Research Fellowship from the National Health and Medical Research Council (NHMRC) Australia, and funds from La Trobe University and La Trobe School of Molecular Science. This work was supported by grants from the NHMRC (APP1160025) to HER and AV, the Cancer Council Victoria Australia (APP1041817) to HER and AV, and the CASS (Contributing to Australian Scholarship and Science) Foundation (SM/13/4847) to LMP, and a National Institute of Health grant GM123136 to AV.

## Author contributions

HER, AV, MP designed the study, MP, LMP, SM, SP, JELM conducted experiments, MP, SM prepared figures, HER, MP wrote the paper, and JELM, AV provided editorial guidance.

