## Supplemental data and figures for "The *Drosophila* Tumour Suppressor Lgl and Vap33 activate the Hippo pathway by a dual mechanism, involving RtGEF/Git/Arf79F and inhibition of the V-ATPase"

**Supplementary Information**

**Supplementary Figure and Legends**

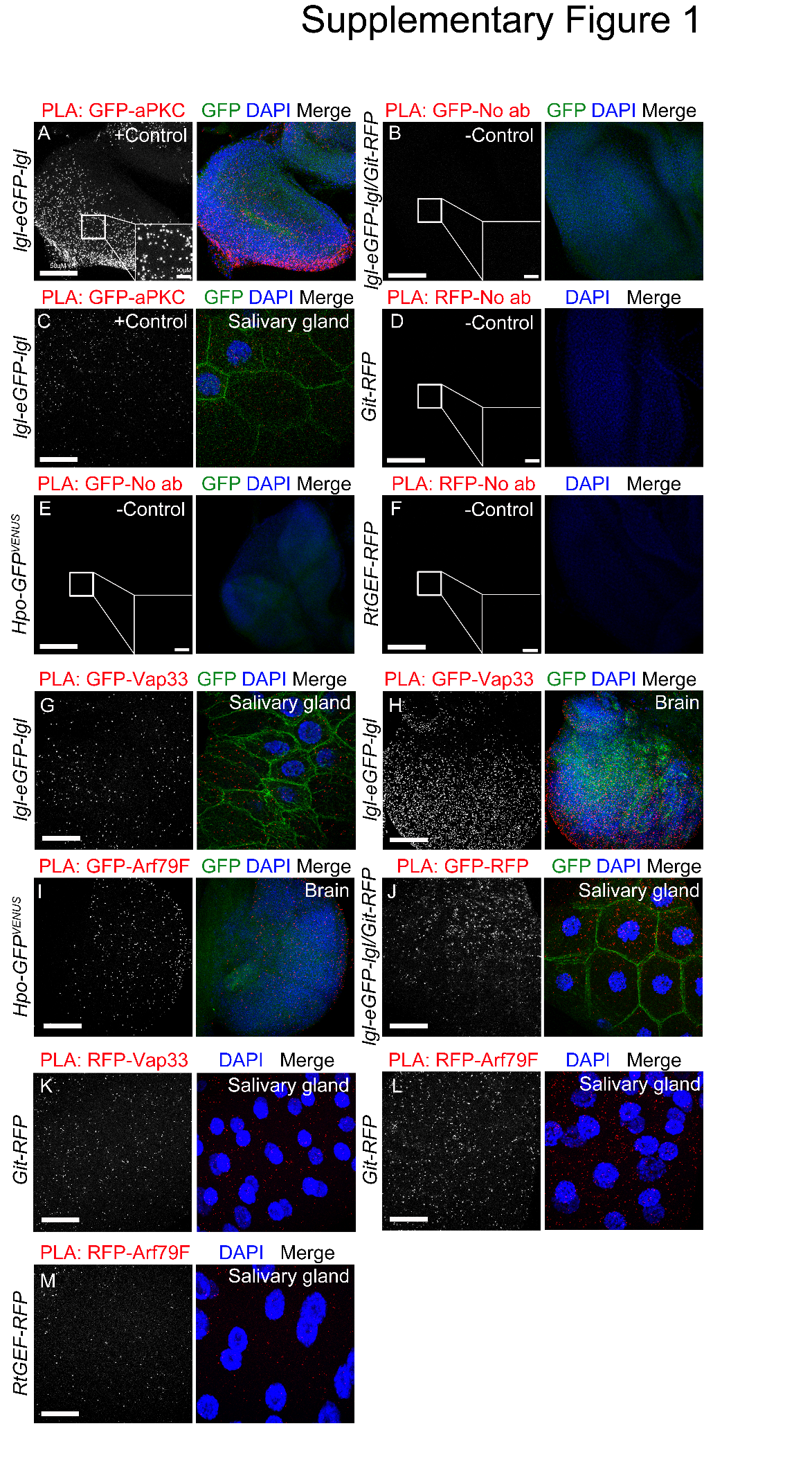

**Fig. S1: RtGEF interacts with Arf79F, Vap33, Lgl and Arf79F interact with Git in the salivary glands, and Arf79F in the brain.**

(A-M) Confocal planar images showing *in situ* proximity ligation assay (PLA) on third instar larval eye discs, brains or salivary glands. Positive PLA results appear as punctate signals (grey, or red in the merges). Nuclei are stained with DAPI (blue). Insets show high magnification images of the PLA foci. (A) Positive-control PLA on *lgl-eGFP-lgl* eye discs using antibodies against GFP and aPKC. (B) Negative control PLA in *lgl-eGFP-lgl*/*Git-RFP* eye discs using only one primary antibody against GFP. (C) Positive-control PLA in *lgl-eGFP-lgl* salivary glands using antibodies against GFP and aPKC. (D) Negative control PLA on *Git-RFP* eye discs using only one primary antibody against RFP. (E) Negative control PLA in *Hpo-GFP* eye discs using only one primary antibody against GFP. (F) Negative control PLA in *RtGEF-RFP* eye discs using only one primary antibody against RFP. (G, H) Positive-control PLA on *lgl-eGFP-lgl* salivary glands (G) and brains (H) using antibodies against GFP and Vap33. (I) PLA in *Hpo-GFP* brains using antibodies against GFP and Arf79F. (J) PLA in *lgl-eGFP-lgl*/*Git-RFP* salivary glands using antibodies against RFP and GFP. (K) PLA in *Git-RFP* salivary glands using antibodies against RFP and Vap33. (L) PLA in *Git-RFP* salivary glands using antibodies against RFP and Arf79F. (M) PLA in *RtGEF-RFP* salivary glands using antibodies against RFP and Arf79F. Scale bars represent 50 μm.

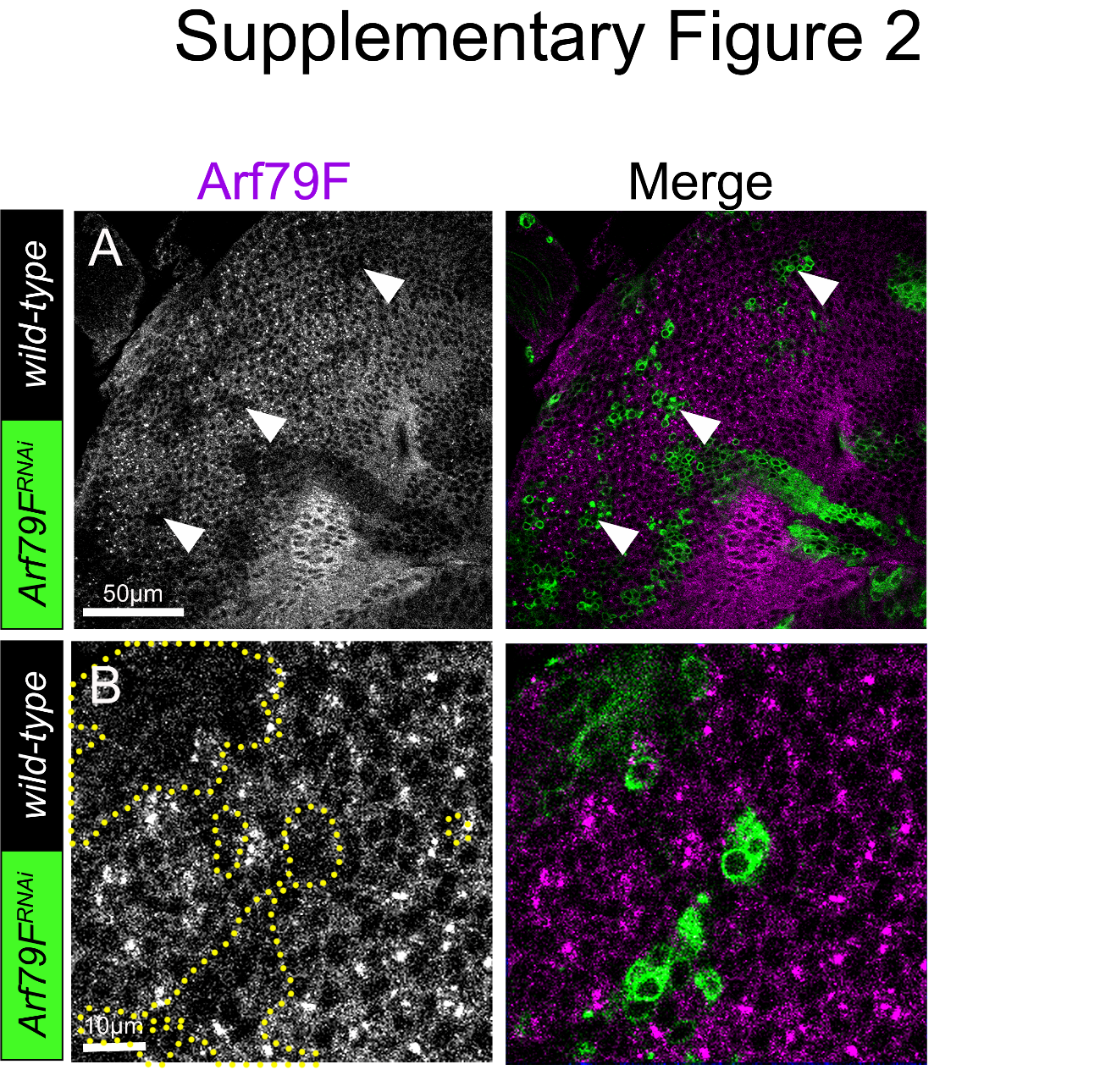

**Fig. S2: Arf79F staining in *Arf79F* knockdown mosaic eye-antennal discs.**

(A) Confocal planar sections of *Arf79F^RNAi^* mosaic third instar larval eye-antennal discs (clones marked by GFP) stained for Arf79F (magenta). Knockdown of Arf79F using the *Arf79F^RNAi^* line reduced the amount of Arf79F protein in the *Arf79F^RNAi^* GFP-positive clones (green, example clones marked by arrowheads). Higher magnification shown in (B). Scale bars represent 50 μm (A) or 10 μm (B).

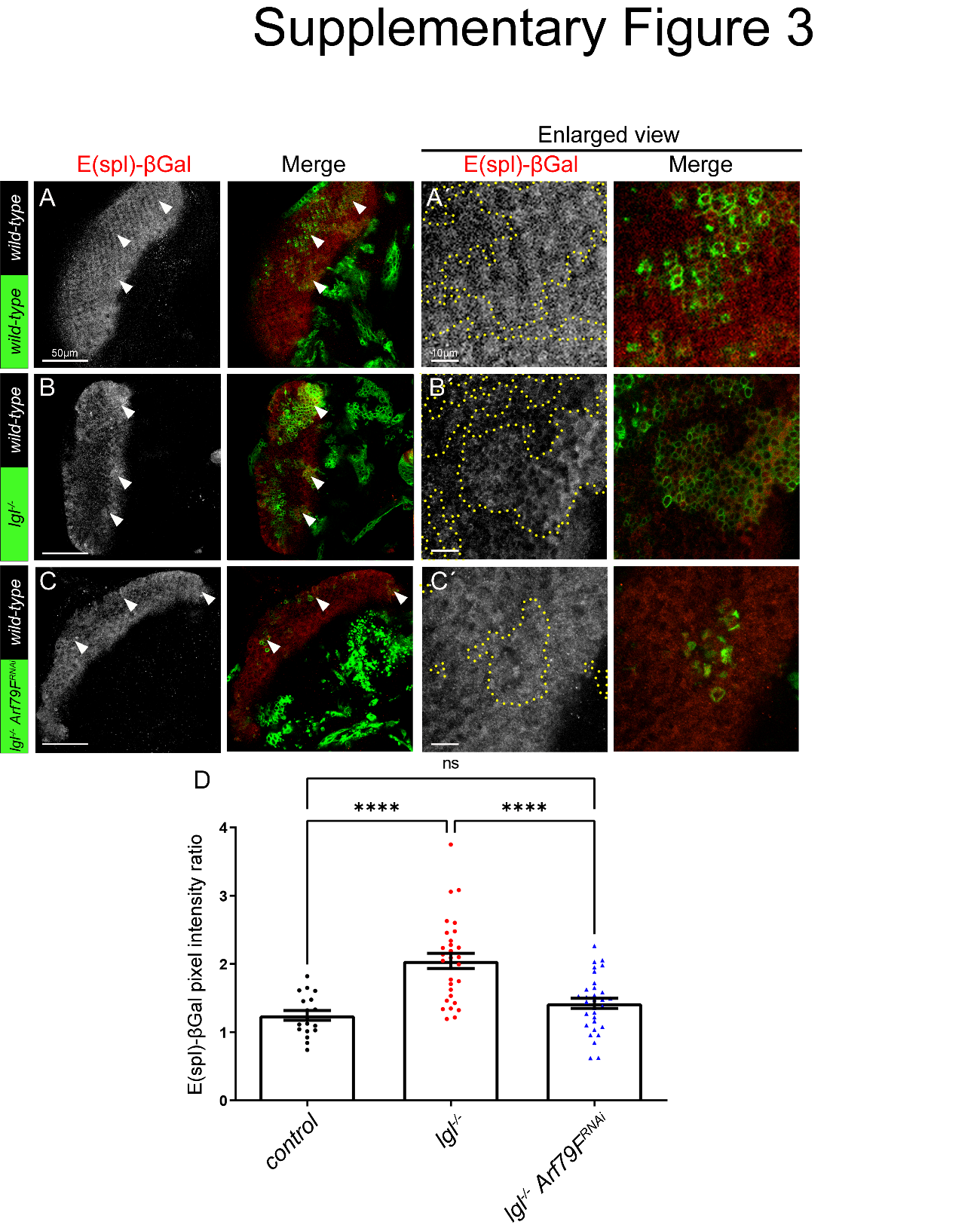

**Fig. S3:** **Knockdown of *Arf79F* prevents Notch signalling pathway upregulation in *lgl* mutant tissue.**

(A-C) Confocal planar sections of a mosaic eye discs containing the Notch target *E(spl)lacZ* reporter, stained for βGal (grey, or red in merges, example clones marked by arrowheads). (A) Control mosaic disc showing endogenous expression of *E(spl)lacZ* within and posterior to the morphogenetic furrow (arrowheads indicate clones showing normal *E(spl)lacZ* expression). (B) *lgl*^-^ mosaic disc (mutant clones are GFP-positive). (C) *lgl*^-^ *Arf79F^RNAi^* (mutant clones are GFP-positive). (D) Quantification of βGal pixel intensity ratio between *wild-type* clones compared to mutant/transgenic clones. Error bars represent SEM. **** P-value<0.0001 (one-way ANOVA with Bonferroni post-test). In all images, posterior is to the left, and the scale bars represent 50 μm. (A´, B´, C´) Higher magnifications of βGal stainings (grey, or red in merges, mutant clones are GFP-positive) for all genotypes. Scale bars represent 10 μm.

**Supplementary Tables**

**Table S1, *Drosophila* stocks**

| ***Drosophila* stock** | **Source and reference** |
| --- | --- |
| *l(2)gl^27S3^* (denoted as *lgl^27S3^*) | [1] |
| *P(E(spl)m8-HLH-2.61)2* (denoted as *E(spl)lacZ^m8-2.61^* or *E(spl)lacZ*,) | A. Bergmann [2] on chromosome *2R* [3] |
| *P(UAS-lacZ.nls)* (denoted as *UAS-lacZ-nls (III))* | G. Baeg |
| *P(UAS-Vha44.P)* (denoted as *UAS-Vha44*) | M. Simons [4] |
| *Vha68-2^R6^* (EMS generated null mutant, due to stop codon mutation) | BL39621, Bloomington Stock Center [5] |
| *P(y[+t7.7] w[+mC]=10XUAS-mCD8::GFP)attP2* (denoted as *UAS-CD8-GFP,*) | BL32184, Bloomington Stock Center |
| *Mi(y[+mDint2]=MIC)l(2)gl[MI07575]* (denoted as *MI07575 lgl-eGFP-lgl* | BL43734, Bloomington Stock Center [6] |
| *P(w[+mC]=UAS-Vap-33-1.P)3* (denoted as *UAS-Vap33*) | BL26693, Bloomington Stock Center [7] |
| *P(TRiP.JF01355)attP2* (denoted as *UAS-luciferase^RNAi^*) | BL31603, Bloomington Stock Center |
| *P(ey3.5-FLP.B)1, y^1^ w^*^; P(w(+mC)=UAS-mCD8::GFP.L)Ptp4E(LL4); P(w(+mC)=tubP-GAL80)LL10 P(ry(+t7.2)=neoFRT)40A; P(w(+mC)=tubP-GAL4)LL7*  (denoted as *ey-FLP, UAS-GFP; Tub-GAL80, FRT40A; Tub-GAL4/TM6B* (*ey-FLP MARCM 2L*)). | [8] |
| *y[1] w[*]; Pvr[1] P(ry[+t7.2]=neoFRT)40A* (denoted as *FRT40A*). | BL58427, Bloomington Stock Center |
| *w; Vha68-2 RNAi TRiP.HMS01056}attP2* | BL34582, Bloomington Stock Center |
| *w;; UAS-Arf79F^RNAi^* | VDRC 23082 and BL66175 |
| *w; UAS-Sec71^DN^* | Fengwei Yu [9] |
| *w; Git-tRFP* | [10] |
| *w; Hpo-VENUS* | [11] |
| *w; RtGEF-tRFP* | [10] |
| *w; FRT40A, RtGEF^1036^/CyO-GFP* | [10] |
| *y v;; UAS-Git^RNAi^* | BL31583 |
| *w;; UAS-wts^RNAi^* | NIG 12072R-1 |
| *ex-lacZ* | BL44248 |
| *yw hs-FLP;; Act>CD2>GAL4, UAS-GFP* | R. Mann |

**Table S2: List of genotypes of the samples used in each Figure.**

| **Figure 1** | **Genotype** |
| --- | --- |
| (A) | *yw hs-FLP; ex-lacZ/ CyO; Act>CD2>GAL4, UAS-GFP/UAS-luciferase^RNAi^* |
| (B) | *yw hs-FLP; ex-lacZ/ CyO; Act>CD2>GAL4, UAS-GFP/UAS-Vha86-2^RNAi^* |
| (C) | *yw hs-FLP; ex-lacZ/ CyO; Act>CD2>GAL4, UAS-GFP/UAS-wts^RNAi^* |
| (D) | *yw hs-FLP; ex-lacZ/ CyO; Act>CD2>GAL4, UAS-GFP/UAS-Vha44* |
| **Figure 2** |  |
| (A-B) | *w, eyFLP, UAS-GFP; lgl27S3, FRT40A; Tub-GAL80, FRT40A, Tub-GAL4* |
| (C-D) | *w, eyFLP, UAS-GFP; FRT40A; UAS-Vap33o/e /Tub-GAL80, FRT40A, Tub-GAL4* |
| (E-F) | *w, eyFLP, UAS-GFP; lgl27S3, FRT40A; UAS-Vap33o/e /Tub-GAL80, FRT40A, Tub-GAL4* |
| **Figure 4** |  |
| (A) | *y,w; MiMIC lgl-eGFP-lgl MI07575* |
| (B) | *w; y,w; MiMIC lgl-eGFP-lgl MI07575*/*Git-RFP* |
| (C, E) | *w; Git-RFP* |
| (D) | *w; Hpo-GFP* |
| (F) | *w; RtGEF-RFP* |
| **Figure S1** | Related to Figure 4 |
| (A, C, G, H) | *y,w; MiMIC lgl-eGFP-lgl MI07575* |
| (B, J) | *y,w; MiMIC lgl-eGFP-lgl MI07575*/*Git-RFP* |
| (D, K, L) | *w; Git-RFP* |
| (E, I) | *w; Hpo-GFP* |
| (F, M) | *w; RtGEF-RFP* |
| **Figure 5** |  |
| (A-B) | *w, eyFLP, UAS-GFP; FRT40A; UAS-Vap33o/e /Tub-GAL80, FRT40A, Tub-GAL4* |
| (C-D) | *w, eyFLP, UAS-GFP; RtGEF^-^, FRT40A; UAS-Vap33o/e /Tub-GAL80, FRT40A, Tub-GAL4* |
| **Figure 6** |  |
| (A-B) | *w, eyFLP, UAS-GFP; FRT40A; Tub-GAL80, FRT40A, Tub-GAL4* |
| (C-D) | *w, eyFLP, UAS-GFP; Vha68-2^-^, FRT40A; Tub-GAL80, FRT40A, Tub-GAL4* |
| (E-F) | *w, eyFLP, UAS-GFP; Vha68-2^-^, FRT40A; UAS-Git^RNAi^ /Tub-GAL80, FRT40A, Tub-GAL4* |
| **Figure 7** |  |
| (A-B) | *w, eyFLP, UAS-GFP; FRT40A; Tub-GAL80, FRT40A, Tub-GAL4* |
| (C-D) | *w, eyFLP, UAS-GFP; lgl27S3, FRT40A; Tub-GAL80, FRT40A, Tub-GAL4* |
| (E-F) | *w, eyFLP, UAS-GFP; FRT40A; UAS-Arf79F^RNAi^ /Tub-GAL80, FRT40A, Tub-GAL4* |
| (G-H) | *w, eyFLP, UAS-GFP; lgl27S3, FRT40A; UAS-Arf79F^RNAi^ /Tub-GAL80, FRT40A, Tub-GAL4* |
| (I-J) | *w, eyFLP, UAS-GFP; FRT40A, UAS-Sec71^DN^; Tub-GAL80, FRT40A, Tub-GAL4* |
| (K-L) | *w, eyFLP, UAS-GFP; lgl27S3, FRT40A, UAS-Sec71^DN^; Tub-GAL80, FRT40A, Tub-GAL4* |
| **Figure S2** | Related to Figure 7 |
| (A-B) | *w, eyFLP, UAS-GFP; FRT40A; UAS-Arf79F^RNAi^ /Tub-GAL80, FRT40A, Tub-GAL4* |
| **Figure S3** | Related to Figure 7 |
| (A) | *w, eyFLP, UAS-GFP; FRT40A, E(spl)m8-lacZ; Tub-GAL80, FRT40A, Tub-GAL4* |
| (B) | *w, eyFLP, UAS-GFP; lgl27S3, FRT40A, E(spl)m8-lacZ; Tub-GAL80, FRT40A, Tub-GAL4* |
| (C) | *w, eyFLP, UAS-GFP; lgl27S3, FRT40A, E(spl)m8-lacZ; UAS-Arf79F^RNAi^ /Tub-GAL80, FRT40A, Tub-GAL4* |
